# Beyond gradients: Factorized, geometric control of interference and generalization

**DOI:** 10.1101/2021.11.19.466943

**Authors:** Daniel N. Scott, Michael J. Frank

**Affiliations:** Department of Neuroscience, Brown University; Department of Cognitive and Psychological Sciences, Brown University; Carney Institute for Brain Science, Brown University

## Abstract

Interference and generalization, which refer to counter-productive and useful interactions between learning episodes, respectively, are poorly understood in biological neural networks. Whereas much previous work has addressed these topics in terms of specialized brain systems, here we investigated how learning rules should impact them. We found that plasticity between groups of neurons can be decomposed into biologically meaningful factors, with factor geometry controlling interference and generalization. We introduce a “coordinated eligibility theory” in which plasticity is determined according to products of these factors, and is subject to surprise-based metaplasticity. This model computes directional derivatives of loss functions, which need not align with task gradients, allowing it to protect networks against catastrophic interference and facilitate generalization. Because the model’s factor structure is closely related to other plasticity rules, and is independent of how feedback is transmitted, it introduces a widely-applicable framework for interpreting supervised, reinforcement-based, and unsupervised plasticity in nervous systems.

## Introduction

When animals learn new skills they often generalize prior learning, and rarely forget or degrade it (Dekker *et al*.2022; Franklin & Frank 2018; Ghirlanda & Enquist 2003; Tenenbaum & Griffiths 2001; Shepard 1987). How biological neural networks support these capacities, including whether and how plasticity rules do, remains poorly understood. For example, previous work has typically addressed interference (memory degradation) and generalization (constructive re-use) using divisions of labor across large scale brain networks (Schapiro *et al*. 2017; O’Reilly & Norman 2002; Mcclelland *et al*. 1995; Frank & Badre 2012; Niv *et al*. 2015; Rougier *et al*. 2005; Collins & Frank 2013; Flesch *et al*. 2021) leaving open the question of how local plasticity rules do or don’t impact such outcomes. Computational work on biological plasticity, by contrast, has often focused on understanding how biology relates to gradient descent (via error backpropagation) in artificial neural networks (Zenke & Ganguli 2018; Bellec *et al*. 2020; Liu *et al*. 2021), or how it might perform unsupervised functions like feature detection (Bienenstock *et al*. 1982). Many empirical studies show, however, that in-vivo plasticity impacts interference and generalization, with cellular and dendritic excitability modulation appearing particularly important in this regard (Cichon & Gan 2015; Yang *et al*. 2014; Sehgal, Filho, *et al*. 2021). We address these sorts of mechanisms theoretically here.

We approached the question of how plasticity impacts interference and generalization (examples in figure 1A-C) by examining neural network loss gradients. We observed that, mathematically, gradients can be decomposed into analogues of population response changes and receptive-field re-weightings, and that interference and generalization are functions of these changes (figure 1D-E). The population responses in this formulation are layer-wise (or “local”) neural response patterns, whereas the receptive fields specify the relative sensitivities of an individual neuron to different input patterns. To make these two concepts more concrete, consider two neurons that initially respond proportionally to a stimulus. If one then increases its firing rate relative to the other, we would refer to this as a change in population response to that stimulus. If, instead, each maintains its response relative to the other but shifts which features drive that response, we would refer to these as purely receptive field changes (for each neuron).

**Figure 1.**
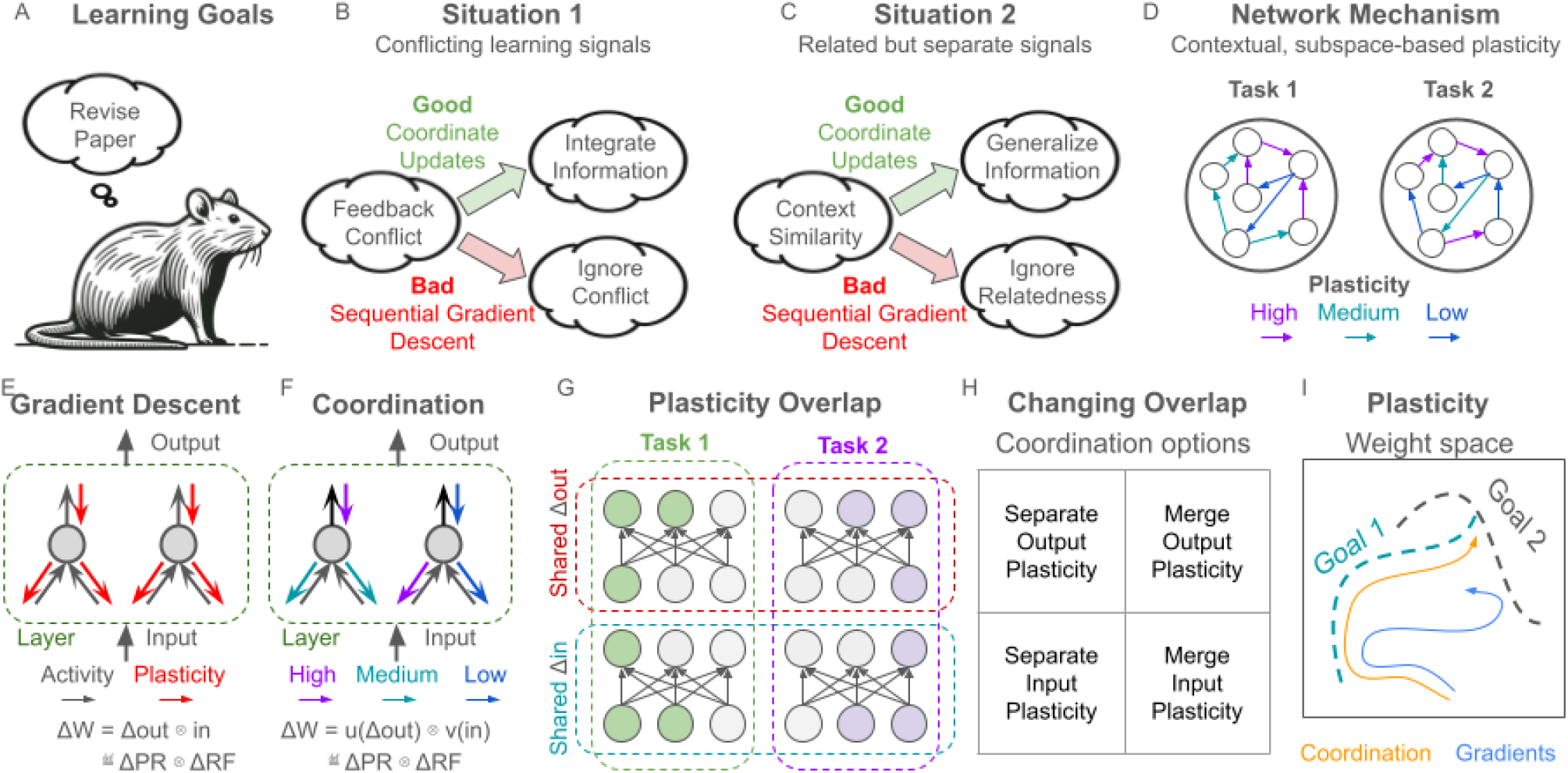
Conceptual overview of this work. (A) Animals have goals, which they must learn to achieve. In this case, consider revising a paper. (B) Some feedback signals (e.g., from reviewers or co-authors) will conflict with one another. An appropriate way to deal with these conflicts is to integrate over learning signals, averaging out noise and clarifying signal (upper arrow). A poor way to deal with conflict is to completely adhere to all feedback, even when it conflicts with other feedback (lower arrow), i.e., to perform sequential gradient descent, which can result in undoing previous learning, rather than reconciling new learning with old. (C) A related situation arises when contextual information suggests generalizing learning. For example, recognizing that two types of feedback reflect the same principle can support learning based on the principle (upper arrow) rather than solely the particulars of the feedback (lower arrow), generalizing learning. (D) Within a network, regularizing plasticity towards particular activity subspaces (which can be shared across contexts) and minimizing the overlap of these subspaces when they interfere, can accomplish these goals. (E) Examining gradient descent, we observe that “input” and “output” or “receptive field” and “population response” factors in a network layer’s plasticity partition this plasticity into subspaces. (F) We explore the idea that independently controlling these two biologically meaningful factors (using functions u and v in the panel) would be useful for avoiding interference and promoting generalization. (G) Example of two tasks that overlap in either RFs (bottom) or PRs(top). (H) We investigate four different scenarios, showing that managing population-response and receptive-field plasticity can avoid interference and promote generalization. (I) These properties result from the fact that coordinating plasticity factors can take arbitrary paths through weight space, whereas gradients always move directly towards individual tasks’ solutions.

Notably, distinct biological processes can control population response and receptive field plasticity and, thereby, network function (for example, Cichon & Gan 2015; Sehgal, Filho, *et al*. 2021). We thus developed a “coordinated eligibility” theory (figure 1D,F), in which structured population-response (PR) and receptive-field (RF) eligibility formed the building blocks of network plasticity. (“Eligibility” refers to the capacity for plasticity, as distinguished from realized plasticity). This theory is closely related to other biological plasticity theories, including three-factor rules (Williams 1992; Fiete & Seung 2006; Izhikevich 2007; Farries & Fairhall 2007; Frémaux, Sprekeler, *et al*. 2013), the BCM model (Bienenstock *et al*. 1982), and two-threshold calcium theories (Evans & Blackwell 2015), supporting its plausibility and making it applicable to understanding these. By exposing tunable parameters of plasticity, which facilitate holistic function, it provides an account of how local plasticity might avoid problems deriving from the memorylessness, and local (rather than global) optimality of gradient descent.

Our mathematical results establish the power and relevance of coordinated eligibility models to in-vivo plasticity, and our simulations provide a further foundation for understanding them. Because we identify two mechanisms for managing between-task learning interactions, and these interactions can be either constructive or destructive, there is a natural set (2×2) of asymptotic network conditions to investigate (figure 1H). Our first four simulations address these cases, using a form of surprise-based metaplasticity to structure population-response and receptive-field changes. In line with our mathematical results, they detail how networks can avoid interference and promote generalization, by projecting population-response or receptive-field changes onto distinct or shared subspaces between tasks. As we show, performing these projections overcomes various limitations of learning strictly via gradient descent, by learning along directional derivatives of task losses (figure 1I). In this context we find that forgetting and incorrect generalization are inversely related, and that this can be understood in terms of the weight geometry of task solutions. Whereas these initial simulations focused on control of either PR or RFs independently, our fifth simulation consider a case in which PR and RF eligibilities are yoked, finding that they allow powerful forms of dimensionality reduction to be embedded in plasticity rules themselves. Together, these results provide an account of interference and generalization in terms of two fundamental biological components, in a framework that can be applied to investigate unsupervised, reinforcement-based, or supervised plasticity in the brain.

## Results

We present our results in a mathematical section and several simulation sections. In the mathematical section we define our basic neural network model and we ask how interference and generalization between tasks are related to weight changes. We find that we can decompose interference and generalization according to “receptive field” (RF) and “population response” (PR) factors of plasticity for each neural network layer. We then formulate our coordinated eligibility theory in terms of these factors. Next, to quantify the impact of coordinated eligibility on task performance, we consider networks which leverage unsupervised metaplasticity to coordinate input and response plasticity components, and we compare them to gradient descent. To do so, we simulate a series of supervised learning problems, aimed at probing the degrees of freedom introduced by the factorized, geometric form of the coordinated eligibility theory. Notably, determining when and where to apply different sorts of metaplasticity is not our goal. Nonetheless, our simulations show how coordinating input plasticity, response plasticity, or both, can effectively avoid interference and promote generalization. Our fifth simulation also initiates work on a fundamentally new direction, that of composing gradient projections according to representational structure across layers, which may facilitate functions such as compositional learning.

### Weight gradients factorize into population-response and receptive-field changes

We consider neural networks performing sequences of tasks. The neural networks are standard multi-layer perceptrons, with layers defined by the equation:

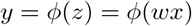

Each layer output *y* is sum of inputs *x* weighted according to matrix *w*, subject to activation function *ϕ*. We define task performance in terms of loss functions denoted *L*. Here we use a supervised learning paradigm, making these losses explicit, but our results apply equally to reinforcement learning and unsupervised settings, where one can often consider learning in terms of implicit loss functions (indeed we demonstrate analogous effects in RL contexts in the appendices, together with a fuller explanation of the application to RL).

The gradient of network parameters with respect to a loss can be computed using the chain rule:

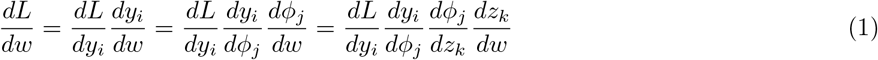

This equation is summed over all indices. Performing some partial sums, as shown explicitly in the appendix, gives an outer-product in terms of a column vector *g* and the layer’s input *x*:

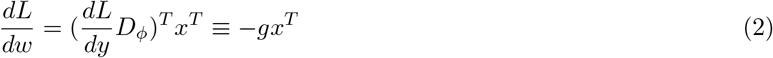

In this equation the column vector *g* summarizes how *y* should change, given input *x*, for the loss to change maximally. The matrix *D*_*ϕ*_ is a diagonal matrix of the local slopes of *ϕ* at each component of *z*. The gradient of the loss with respect to *y* (the term *dL/dy*) captures how downstream processing makes use of the local post-synaptic firing rates, ultimately manifesting in a loss value. In the reinforcement learning and unsupervised cases, this *g* is typically accumulated through experience, whereas supervision and backpropagation implicitly determine it. This representation of the gradient is striking in relation to neuroscience, because the *x* vector denotes how a given neuron’s inputs should be re-weighted (i.e., plasticity of dendritic processes and receptive fields), whereas the *g* vector denotes how a layer of neurons’ collective output is changed (i.e., plasticity of population response patterns). In particular, this parallels the fact that individual neurons can independently exhibit both gain (or excitability) changes and re-weighting of their inputs (see, for example, Cichon & Gan 2015; Yang *et al*. 2014; Sehgal, Filho, *et al*. 2021; Scott & Frank 2022). As such, processing can change when population responses change, inputs are re-weighted with respect to one-another, or both. For example, a weight change Δ*w* = *uv*^*T*^, can involve input reweighting (changes in the row space of *w* arising from *v*), population response changes (differences in the column space of *w* arising from *u*) or both. We hypothesized that this would allow interference and generalization between tasks to be managed at these two distinct loci, and we show that this is the case below.

### Gradient factors determine interference and generalization

For a weight change to impact performance, by definition, it must have a non-zero inner product with the gradient of a loss. Indeed, the loss gradient is defined to be the direction in weight space along which that loss increases maximally. Hence an orthogonal change in weights (i.e., in the loss’ null-space) will neither increase nor decrease the loss. If we consider two rank-1 weight changes that are outer products (such as gradients for two tasks), denoted 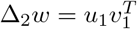 and 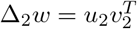, then these have an inner product:

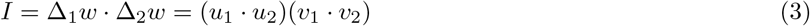

This equation shows that both the input and population response components of plasticity must overlap for interference or generalization to occur. Intuitively, gradient updates interfere with one another when they change network responses to the same input in opposite ways, for example when *u*_1_ = *g*_1_ = −*g*_2_ = *u*_2_ and *v*_1_ = *x*_1_ = *x*_2_ = *v*_2_. Updates generalize across tasks when they change the network in similar ways, such as when *u*_1_ = *g*_1_ ∝ *g*_2_ = *u*_2_ and *v*_1_ = *x*_1_ = *x*_2_ = *v*_2_. Both situations are illustrated in figure 2A, using the minimal example of trying to learn two conflicting (non-proportional) input-output mappings using one input and one output neuron (see the caption for further details). In multi-dimensional networks, interference occurs when weights are changed in opposite directions by receptive field plasticity, population response plasticity, or both. These situations are illustrated in figure 2B and 2C, and described further in the caption. Note, however, that if a network only changes responses to two completely distinct stimuli, as when *v*_1_ · *v*_2_ = *x*_1_ · *x*_2_ = 0, then no interference occurs. The same is true when the stimuli themselves are identical, but their processing is completely segregated.

**Figure 2.**
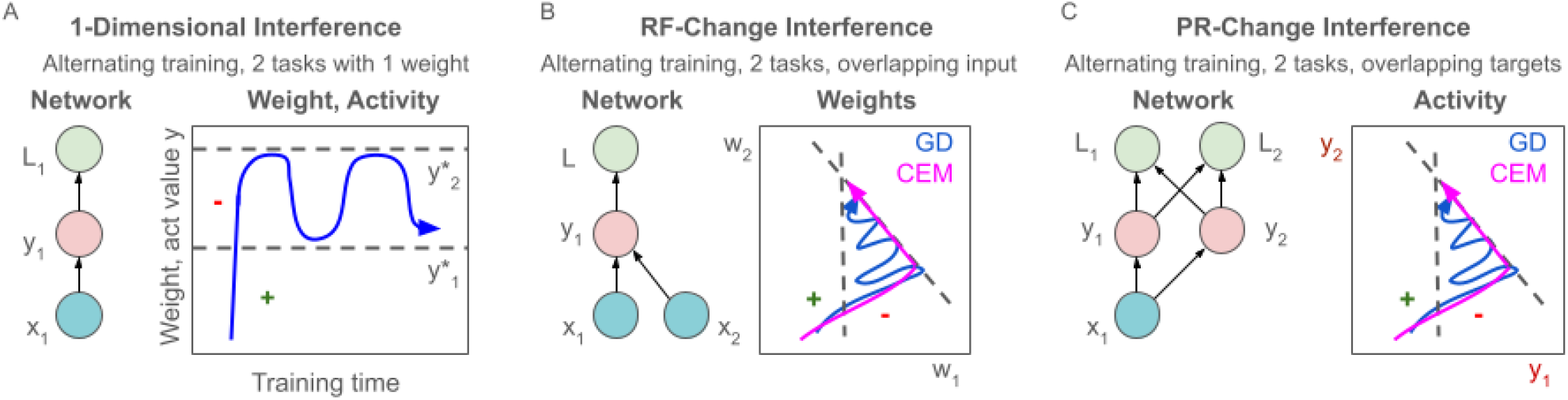
Illustrations of interference from PR and RF plasticity. (A) A simple linear network, with one input neuron (x), one response neuron (y), and a loss function (L). If an input *x* has a target output *y*^∗^(1) for one task, and *y*^∗^(2) for a second, then alternating training will pull the weight connecting *x* and *y* in opposite directions. Notice that if the output activity started on one side of both *y*^∗^(1) and *y*^∗^(2), then there would first be a period of generalization, during which performance improved on both tasks while training either one. In the lower panel, showing weight/activity dynamics over time, these regions of the weight/activity space are denoted with + and -symbols indicating generalization and interference, and the task-solutions are denoted with dashed lines. The basic illustration is also representative of higher dimensional cases; in more complex networks the main question is how these phenomena are distributed over groups of neurons and weights, instead of individual ones, and multiple tasks, instead of task pairs. (B) A network with two inputs, illustrating RF-change induced interference. Here, training changes weights in two dimensions generating a pattern of regions of interference and generalization based on current weights and the angles between task solutions. Lower left panel:Gradient descent is subject to the same oscillatory behavior in the interference-producing region between task solutions. Lower right panel:By restricting weight update dimensions, one can avoid interference. (C) A network with two outputs, illustrating PR-change induced interference, analogous to B.

### Coordinating eligibility controls learning interactions

Observing the facts above about gradients, interference, and generalization, along with the many biological plasticity controllers, we formalize a coordinated eligibility theory of plasticity. Note that “eligibility” refers to the capacity for plasticity, whereas plasticity refers to synaptic weight (and hence network) change itself. In this theory, plasticity takes the form:

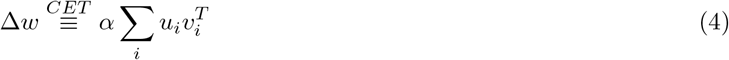

Here, *u* and *v* are generic vectors, *α* is a scalar, and *i* is an index with an arbitrary range.

Many specific forms of plasticity are instances of equation (4). For example, when there are as many terms *i* as synapses, and *u*_*i*_ and *v*_*i*_ are arbitrary, we can capture every possible weight update. When *i* = 1, *u* = −*g* and *v* = *x* we recover gradient descent. Other choices interpolate between these two extremes of flexibility, and other canonical models can also be described using equation (4). Setting *i* = 1, 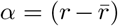 (with *r* denoting “reward”), 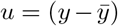, and *v* = *x* gives a REINFORCE algorithm (Williams 1992), which can approximate gradient descent by estimating *g* over time. Specifically, in this latter case, population response variability drives sample-based integration of the gradient term (*α* and 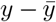 combine in expectation to form *g*). By generalizing this classic REINFORCE model to use non-isotropic sampling (Cov(Δ*y*) ≠ *I*), one arrives at a coordinated eligibility model that effectively projects *g*, which can have much lower sample complexity, along with some of the geometric properties we discuss here.

More broadly, a reasonable set of models to consider might take *α, u*, and *v*, to be functions of *g* (without necessarily requiring *g* to be known explicitly) and *x*, the weight update to have many fewer degrees of freedom than *n*× *m* for *w* ∈*R*^*n×m*^, and the update itself to be distinct from the loss gradient of some task at hand. We investigate several such models below. Specifically, in our simulations we take *α* to be 1, and the factors *u* and *v* to be functions of the gradient components *g* and *x*. These gradient components are projected onto different subspaces according to task demands by matrices *P* and *Q*, and normalized (projected onto the unit sphere via transform *S*). Mathematically:

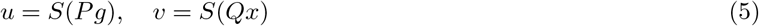

This is one of the simplest coordinated eligibility models, but is flexible enough to avoid interference and promote generalization. Specifically, neglecting some technicality related to normalization, the inner product between two weight updates (with subscripts 1 and 2) will then be:

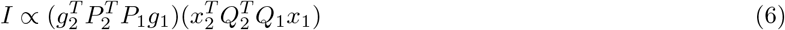

The column spaces (or, roughly, outputs) of the transformation matrices can therefore be used to increase or decrease either quantity in parentheses.

There are many ways these transformations could be manipulated biologically, but we observe that simple, surprise-based, unsupervised metaplasticity can manage interference and generalization. (Both unsupervised and surprised based metaplasticity are well established empirically. See, for example, Yagishita *et al*. 2014; Jaskir & Frank 2021; Scott & Frank 2022) In our simulations below, networks compute surprise over inputs and feedback signals, in order to determine when and how to change the transforms *P* and *Q*. This allows them to perform pattern-separation (in the high-surprise case) or integration-based coarse-coding (in the low-surprise case) of plasticity components *u* and *v*, by progressively removing dimensions from the column spaces of *P* and *Q*.

### Pattern-separating receptive field plasticity reduces interference

To quantify the impacts of coordinating eligibility, we first simulated one of the simplest continual learning problems, online linear regression. Simulating a linear regression problem had the advantages of allowing us to use an easy-to-understand linear neural network, and therefore to compare the network’s solution at each time to the true (minimum norm) solution in terms of both the network’s function and its weights. We demonstrate below that our coordinated eligibility model converges to the true optimal solution for the whole curriculum, whereas gradient descent does not (despite learning locally optimal solutions for each task, sequentially). By comparing plasticity models, this analysis also reveals important properties of both learning algorithms.

Our linear regression problem was composed of 80 input-output associations (“tasks”) with 100-dimensional unit-normal input vectors and 20-dimensional unit-normal target vectors, drawn at uniform angles (figure 3A-3B). These pairs were learned completely and sequentially in one sweep through the data (figure 3C), using either gradient descent or the coordinated eligibility model. A sweep was defined as reduction of the RMSE for a task from an initial value of 1 to a criterion value of 0.05. Mathematically, this resulted in neural networks approximately solving the ordinary least squares regression equations:

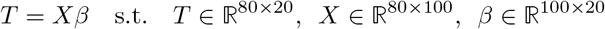

**Figure 3.**
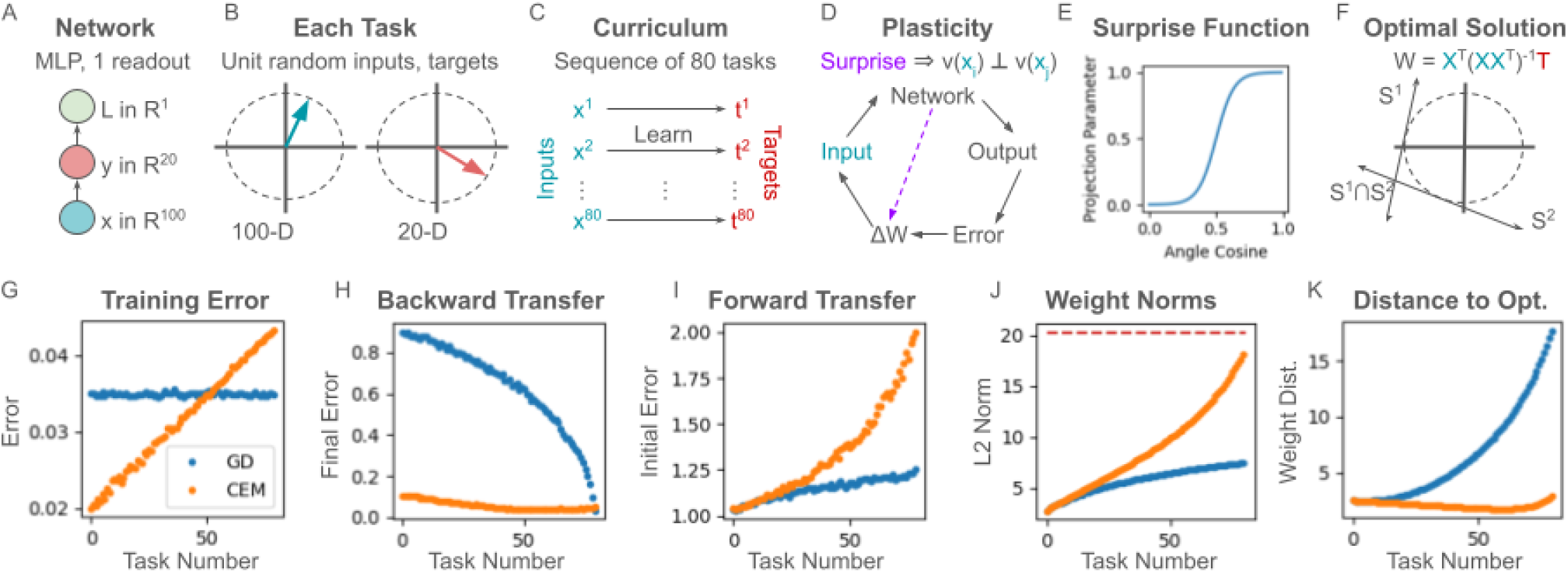
Receptive-field eligibility separation. (A) The network we used was single linear perceptron layer, with a single readout. (B) Tasks in this simulation were each defined by random, unit-normal, 100 dimensional input vectors and similarly distributed 20 dimensional target vectors. (C) Training was performed sequentially over 80 such tasks. When outputs were within 0.05 units of Euclidean distance of targets, training proceeded to the next task. (D) Networks computed surprise over inputs, which was used to determine task change-points and orthogonalize new input plasticity vectors against previous plasticity. (E) The surprise function used was a logistic curve over input cosine angle. (F) The optimal set of weights for the curriculum was computed, for comparison with network outputs, using the pseudo-inverse of the inputs. Intuitively, the solution is the intersection of individual task solutions, which themselves are rank-1 outer products between the (unit normal) inputs and targets, shown here as lines intersecting a unit sphere. (G) Error on each task, computed after training that task. (H) Backward transfer on each task, i.e. task errors at the end of curriculum training. (I) Initial task error on new tasks at each point in curriculum learning (forward transfer). The CEM shows negative transfer related to the fact that it remembers previous inputs, whereas this is reduced, but still present, for GD. (J) Layer weight norms in both models, over the course of learning. CEM weight norms grew over the course of learning to match the optimal network weight norm, given by the dashed red line, whereas GD does not. GD struggles to leave a region of weight space proximal to all individual task solutions, but not their intersection. (I) Distance from the optimal set of weights, indicating that not only is the weight norm of the CEM solution growing properly, the network is also converging to the optimum rather than diverging in an inappropriate direction. By contrast, GD gets further from the curriculum solution over time.

The minimal norm solution (and hence the optimal network weight matrix) is given via a multiplication of T (a matrix of target vectors) by the pseudo-inverse of *X* (a matrix of input vectors):

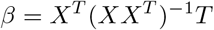

This equation is appropriate regardless of the number of tasks (columns of T and X) the network has seen, and therefore defines an optimal solution after every individual task is learned.

To solve this problem, we simulated a single-headed, single-layer linear network with a squared-error loss. (We also obtain similar results in reinforcement learning simulations using categorical action choice, cross-entropy loss, and nonlinear activation functions, however, which we discuss in the appendices.) In this setting the transformation from weights to a loss value has a large null space, meaning many weights are compatible with solving the task (which is why the weight matrix is a pseudo-inverse rather than an inverse). As a result, upon learning the first input-output pair, the weights “listening to” every other input dimension are free to vary, because the first input has no projection along these. Each of these additional dimensions is constrained only once it is associated with an input, and a learning algorithm can therefore proceed by “listening” only to unique dimensions in new inputs (learning with orthogonal input plasticity vectors *u*) to avoid interference. Given these observations, we set the network’s surprise-based metaplasticity to detect input change-points (using the surprise function in figure 3E), and to progressively remove dimensions of input plasticity when these change-points occurred (figure 3D). How and when networks would know to use this exact form of metaplasticity is an interesting open question, but we introduce it here to demonstrate our points about the geometry of our plasticity factors. We expected this geometry to produce a path toward the optimal solution in weight space that would stay within the solution manifolds identified for previously learned tasks (figure 3F).

We compared the properties of our coordinated eligibility model with gradient descent by examining training error, forward and backward transfer, and network weights, averaged over 200 simulations (figure 3G-K). Training error at the end of training for each task was consistently just below criterion, per expectations, with the margin decreasing over task number, consistent with networks taking steps which were less direct relative to gradients as fewer dimensions of the input space became available for learning (figure 3G). Backward transfer was assessed by testing each task after training the entire curriculum. Networks using gradient descent completely unlearned task 1 by this time, and showed graded forgetting as a function of time-since training for each task (figure 3F). By contrast, the coordinated eligibility model showed minimal interference (figure 3H). We also observed that both gradient descent and the CEM introduced significant negative forward transfer (figure 3I). That is, solving tasks made the initial, untrained error on new tasks larger. This is expected, because each task defines a solution space of weight matrices, and the intersection of a set of these task solution spaces gets further from the origin as their number increases. Because the tasks were normalized, with unit-normal inputs and targets, predicting the 0 vector for new inputs will result in an error of 1, and the further from the origin new untrained task outputs are, the larger the initial error is likely to be. This led us to suspect that gradient-descent was under-fitting the data, and remained in the vicinity of the origin by virtue of moving directly towards each new task’s individual solution subspace.

To test this hypothesis about the nature of the forward transfer problem, we computed the norm of the network’s weights after completing every task, and similarly checked the distance from (curriculum) optimality at each point. We observed that the weight norms under gradient descent were consistently low, relative to the norm of the full solution, whereas the CEM weight norms grew to match this norm (figure 3J). Intuitively, this resulted in the gradient-based solution becoming more distant from the optimal solution as more tasks were completed (figure 3K).

### Pattern-separating population response plasticity reduces interference

Because of the symmetry of relationship (3) between population-response and receptive-field changes, interference and generalization can also be manipulated by changing population responses. Indeed, there are some data suggesting this may occur in the brain, with memories being increasingly segregated according to their distance in time as a result of (for example, CREB-based) excitability drift (Sehgal, Zhou, *et al*. 2018; Lisman *et al*. 2018). Here, we model a complementary (and analogous) situation to the previous simulation, involving population-response based pattern separation, using task-dependent readouts. We describe a pure form of this scenario below, in which interference can only be managed via population response plasticity, because network inputs don’t change. As such, our simulation is a model of purely contextual behavior, with context inferred based on network history.

Specifically, we simulate a continual learning scenario in which the input is constant, but target outputs change across tasks. To do so, we used a single linear layer with a constant scalar input and 100 response neurons (figure 4A). The response neurons were subject to a sequence of unit-random, 100 dimensional readout weights, one defined for each subsequent task, of which there were 80. Without loss of generality, each readout had a target response of 1, defining a 100 dimensional target at the layer output (figure 4B). Networks received each input, were trained to produce the paired target (up to an SSE of 0.05), and then were trained on subsequent input-target pairs, until the whole curriculum had been seen (4C). Mathematically, jointly solving all tasks in the curriculum is equivalent to solving the following equation for *w*:

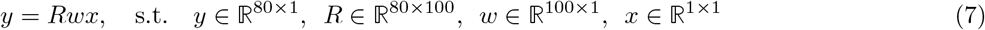

**Figure 4.**
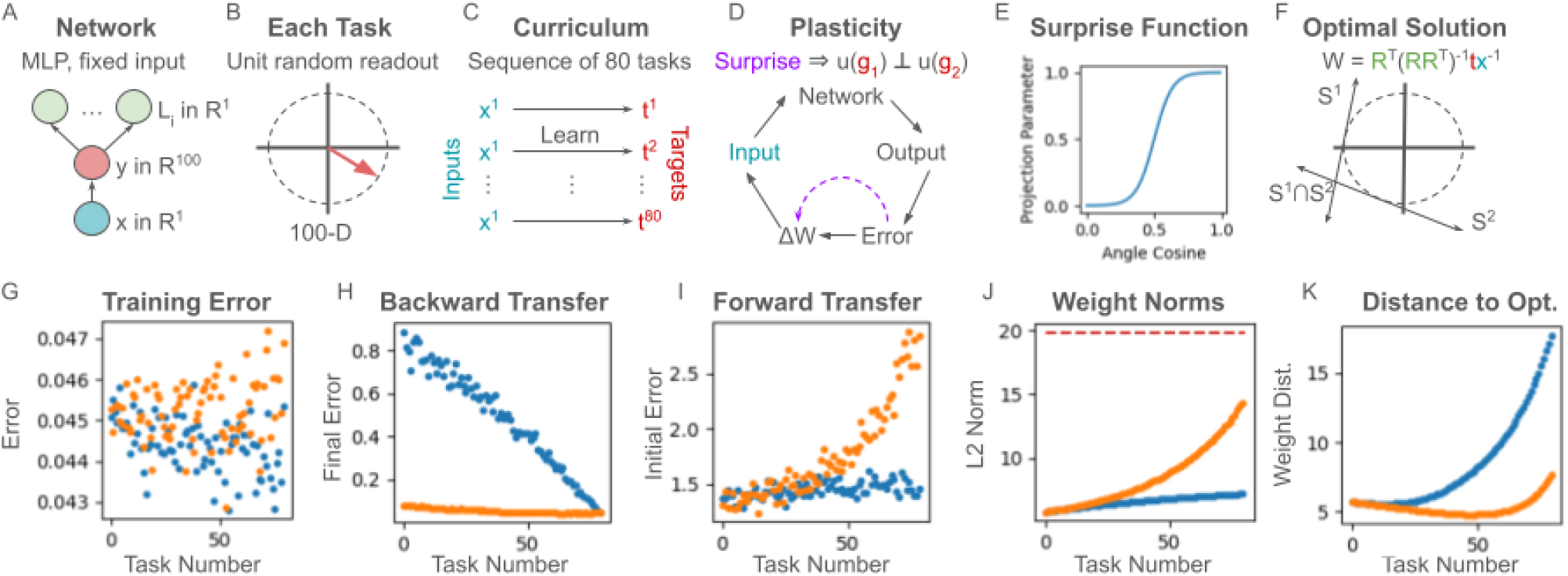
Population-response eligibility separation. (A) These simulations used linear network layers subject to linear readouts, one for each task. (B) Each task is defined by a new random 100-dimensional readout. (C) Tasks are solved sequentially, to within a small error bound, as with the previous simulation, before a new random readout is drawn and applied to initiate learning a new task. (D) Updates to firing-rate response subspaces are orthogonalized based on surprise computed over feedback (gradient) components. (E) Surprise is computed as previously, using a logistic function over relative angles. (F) Optimal weights are analogous to those of the previous simulation. (G) Training error both networks completely solve each task in the curriculum. (H) Gradient descent shows significant forgetting, whereas the CEM does not. (I) Remembering earlier learning produces negative forward transfer, as previously. (J) GD fails to push weights outside the region around the origin, causing them to (K) become increasingly far from optimal as new tasks are seen.

As with the simulation above, this is a standard linear regression problem. Applying a continual learning algorithm will therefore approximate the solution:

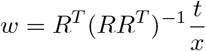

To remove interference in this setting, networks computed surprise over feedback change-points (figure 4D), using a logistic curve over output-plasticity cosine similarity (i.e., in a manner analogous to the previous simulation) (figure 4E). As with the previous simulation, this surprise signal was used to orthogonalize ongoing plasticity from previous plasticity, following these change-points.

After training networks to solve this sequence of tasks, we observe that backward transfer again completely abolishes earlier learning under gradient descent (figure 4H). Here, too, we have averaged over simulations (400), and again, the coordinated eligibility model does not suffer from negative backward transfer (figure 4H). Forward transfer results are also analogous to the previous simulation (figure 4I), and appear to be explained by the same difference in the algorithms’ abilities to effectively leave the origin of the networks’ weight spaces (figure 4J-K). As with our previous simulations, we note that we obtained similar results in reinforcement learning simulations (also discussed in the appendices).

### Coarse-coding receptive field plasticity generalizes learning

A network that perfectly pattern separates all learning will completely avoid interference, at the potential cost of also removing positive transfer between tasks (i.e., generalization). By contrast, a network that coarse-codes or generalizes receptive field plasticity will produce generalized learning, while potentially risking negative transfer. In any given setting, there are therefore sets of desirable basis functions for receptive-field plasticity, and sets of undesirable ones. In the previous section, we addressed the removal of undesirable generalization (i.e., interference) by restricting plasticity, and in this section, we explore the promotion of desirable generalization via coarse-coding. In the pattern-separation simulations above, our network had a built-in assumption that inputs with sufficiently different angles represented different tasks, defining boundaries across which learning should be separated. However, there are many complementary situations in which superficially dissimilar states should instead be considered equivalent in a more abstract sense. This is typically studied in the context of state abstraction and task-set clustering across contexts in reinforcement learning Lehnert *et al*. 2020; Collins & Frank 2013, whereby such abstraction is needed for generalization.

To motivate this situation, consider the case of object affordances. Many classes of objects are loosely defined by their uses, such that (for example) the category of mugs is a set of things one drinks from. Such uses create natural categories for generalizing learning. If we interpret a target network output as an action, use, or label, then the inputs which should prompt a given output can often be inferred by learning the input transformations which the target output is invariant to. For example, if I see several distinct mugs, I can note that those physical transformations taking one mug to another should generally leave their shared property of being open-top vessels intact; this will allow me to drink out of the transformed objects. Knowing this set of transformations will increase my ability to say which other sorts of objects might be mugs, and therefore to generalize inferences or associations over this class. In this case, learning (plasticity) could be coarse-coded by yoking receptive-field changes *v* (which represent changes to how mug features are detected) for a second labeling task over category members from the first (i.e., according to similarity over vectors *u*, representing the mug-category response). For example, by observing that I can’t drink coffee out of a particular floppy mug, I may be justified in learning that “can be floppy” is generally false for open-top-drinking-vessels broadly, and therefore other mugs specifically. A similar plasticity-component logic would apply in other generalization scenarios, and structurally equivalent forms of generalization have been studied empirically in state abstraction and task-set clustering situations Collins & Frank 2016).

In analogy with this example, we simulated a continual learning problem in which object classes had (by definition) the same target outputs, and were seen in blocks. During the first block, networks learned one distinct target output for each group of inputs. We refer to this first block as an association block. Target similarity, computed as cosine angle, was then used to induce a similarity measure on inputs, which itself defined coarse-coded input plasticity in a subsequent block of trials (a generalization block). In this second block of trials, we model a new group of tasks using a separate regression head, and demonstrate that the coordinated eligibility model is able to effectively transfer learning within classes defined by the first block.

Concretely, we simulated two linear regression problems, using a network with 100 dimensional inputs, 20 dimensional outputs, and two readouts (figure 5A). All of these were random unit vectors (figure 5B). Inputs during the association task (“object exemplars”) were randomly paired (and hence had no intrinsic similarity), and pairs had the same associated target (“object property or class”). During the association curriculum, networks were trained to produce the correct target for each object, sequentially (figure 5C). Surprise was computed over targets (technically, population response feedback) to establish class-presentation change-points. Future receptive field plasticity was then coarse-coded over the inputs that had been seen during the previous low-surprise period (figure 5D). Subsequently, during a generalization task, new targets were generated for each existing pair of inputs, on a new readout (i.e., old inputs/classes were conserved) (figure 5E). These new input-target associations were trained (sequentially, in a curriculum) on one of each pair of inputs and tested on the other (figure 5F). Performance was assessed at each stage of the task (figure 5G), verifying that both gradient descent and the coordinated eligibility model performed identically at initialization, after association training, and after training during the generalization phase. The algorithms differed in terms of their generalization however (by construction), with coarse-coded plasticity ensuring that held-out inputs were properly associated with trained targets during the generalization phase.

**Figure 5.**
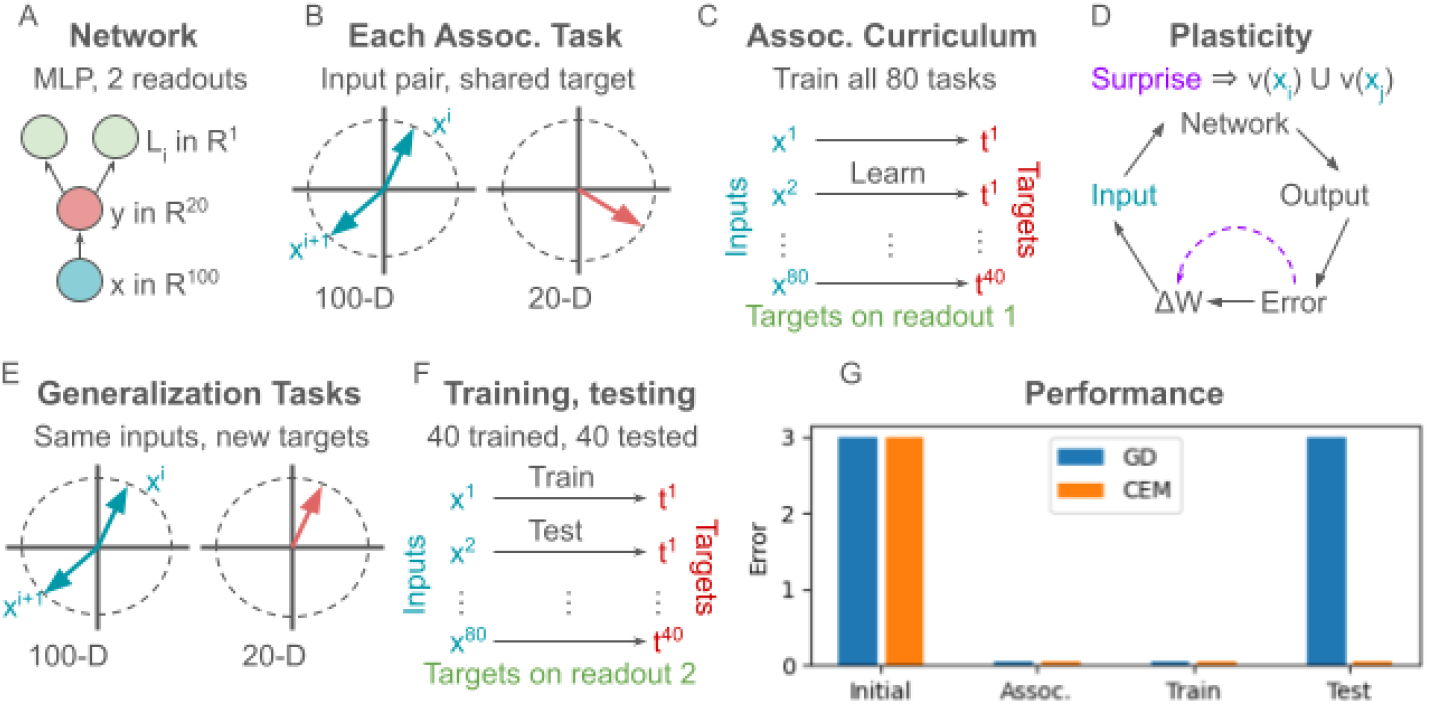
Receptive field pattern completion. (A) The network we used was an MLP with one readout (head) for the association curriculum and one for the generalization curriculum. (B) Inputs were 100 dim. unit random vectors. Targets were 20 dim. unit random vectors. (C) During the association phase, the network learned to map pairs of inputs, presented sequentially, to targets (of which there was one for each pair). (D) Surprise was computed over target prediction errors and used to chunk learning temporally, producing a coarse-code for gradient components between elevated surprise events. (E) A generalization curriculum re-used inputs from the association curriculum, but paired them with new targets. (F) Training was performed on only one item out of each pair, testing was performed on the held out item subject to coarse-coded plasticity. (G) Initial error, error at the end of the association-learning phase, error at the end of training during the generalization phase (for trained items) and generalization (test) error at the end of the generalization phase. Coarse coding generalized learning.

### Coarse-coding population response plasticity generalizes learning

As with pattern separation, pattern completion (or coarse-coding) can also be applied over population responses. We illustrate this by simulating a network performing tasks analogous to the RF pattern completion case just discussed, with PR and RF roles reversed. Specifically, we first simulated a curriculum in which pairs of targets (on independent readouts) were associated (via training) with individual inputs (figures 6A-C). At every input transition, input-based surprise was used to coarse-code plasticity for the previously seen targets (which shared the previous input) (figure 6D). As previously, networks made use of this coarse-coding during training on a generalization curriculum (figure 6E-f). During these tasks, networks were given a sequence of new inputs, and were required to produce one out of of each of 40 pairs of targets. Plasticity during training was constrained to the coarse-coded dimensions discovered previously. Testing the network on the held-out targets verified that it properly produced each second target, given each held-out readout, and that performance at other phases of learning was otherwise identical (figure 6G).

**Figure 6.**
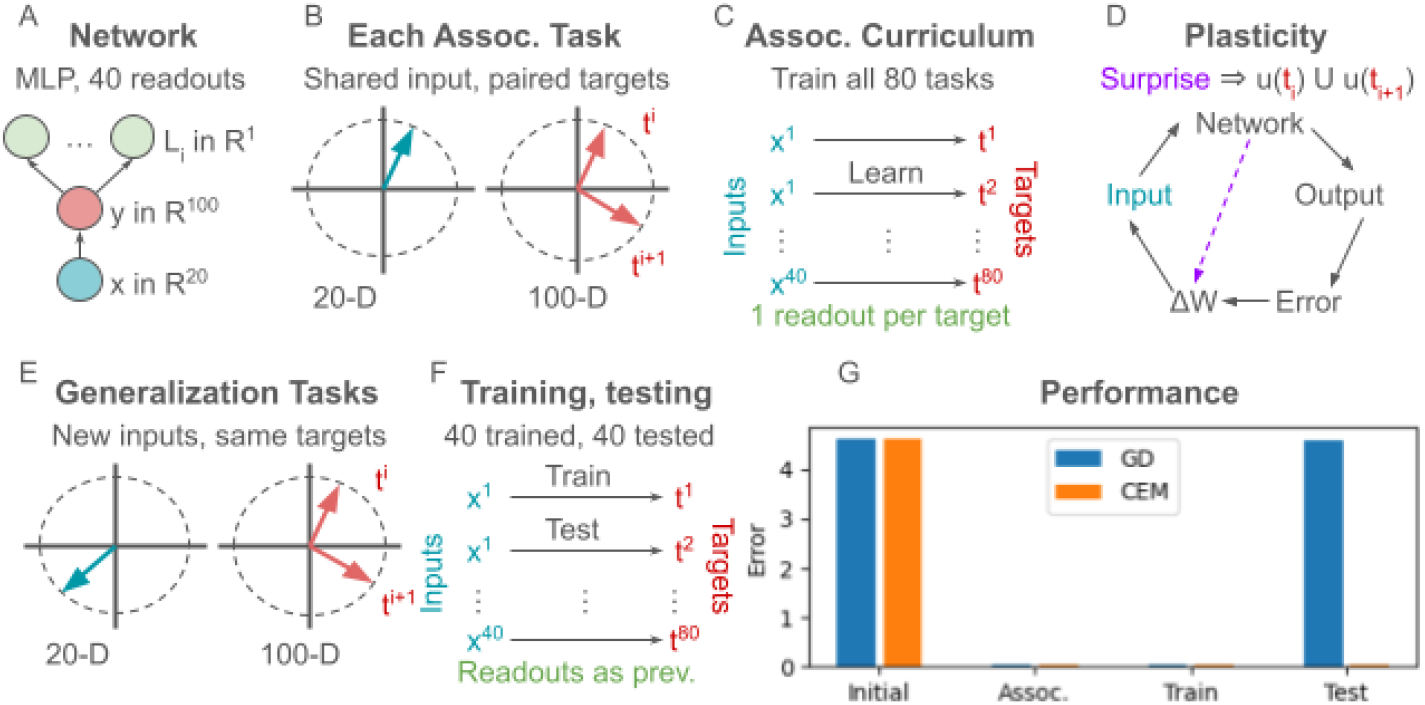
Population response pattern completion. (A) The network we used was an MLP with one readout (head) per target in the association curriculum, and these were re-used for the generalization curriculum. (B) Inputs were 20 dim. unit random vectors. Targets were 100 dim. unit random vectors. (C) During the association phase, the network learned to map inputs, presented sequentially, to pairs of targets (one pair per input). (D) Surprise was computed over inputs and used to chunk learning temporally and coarse-code gradient components between elevated surprise events. (E) A generalization curriculum re-used readouts from the association curriculum, but paired them with new inputs. (F) Training was performed on only one target out of each pair associated with a given input, while testing was performed on the held out readout, subject to coarse-coded plasticity. (G) Initial error, error at the end of the association-learning phase, error at the end of training during the generalization phase (for trained targets) and generalization (test) error at the end of the generalization phase. Coarse coding generalized learning.

### Eligibility components are compositional

Moving beyond simple subspace restrictions in our coordinated eligibility theory, we examined tasks with compositional inputs, requiring compositional responses. Many objects can be described as bundles of features of varying statistical interdependence, and when learning a task, subsets of these features might dictate separate elements of an appropriate response (figure 7B). Brain regions with convergent input from diverse pre-synaptic partners would be well served by plasticity mechanisms that could manage responsiveness to the compositional building blocks in such mixed inputs, and the resulting compositionality would also facilitate combinatorial generalization (as in e.g. O’reilly 2001). While there are circuit level architectural mechanisms that can gate attention to specific features in order to govern responding (Frank & Badre 2012; Niv *et al*. 2015), just as there are for pattern separation vs completion O’Reilly & Norman 2002, we again consider here how learning rules themselves can implement compositional inductive biases. As we show, associations between population-response plasticity and receptive field plasticity can produce compositional learning.

**Figure 7.**
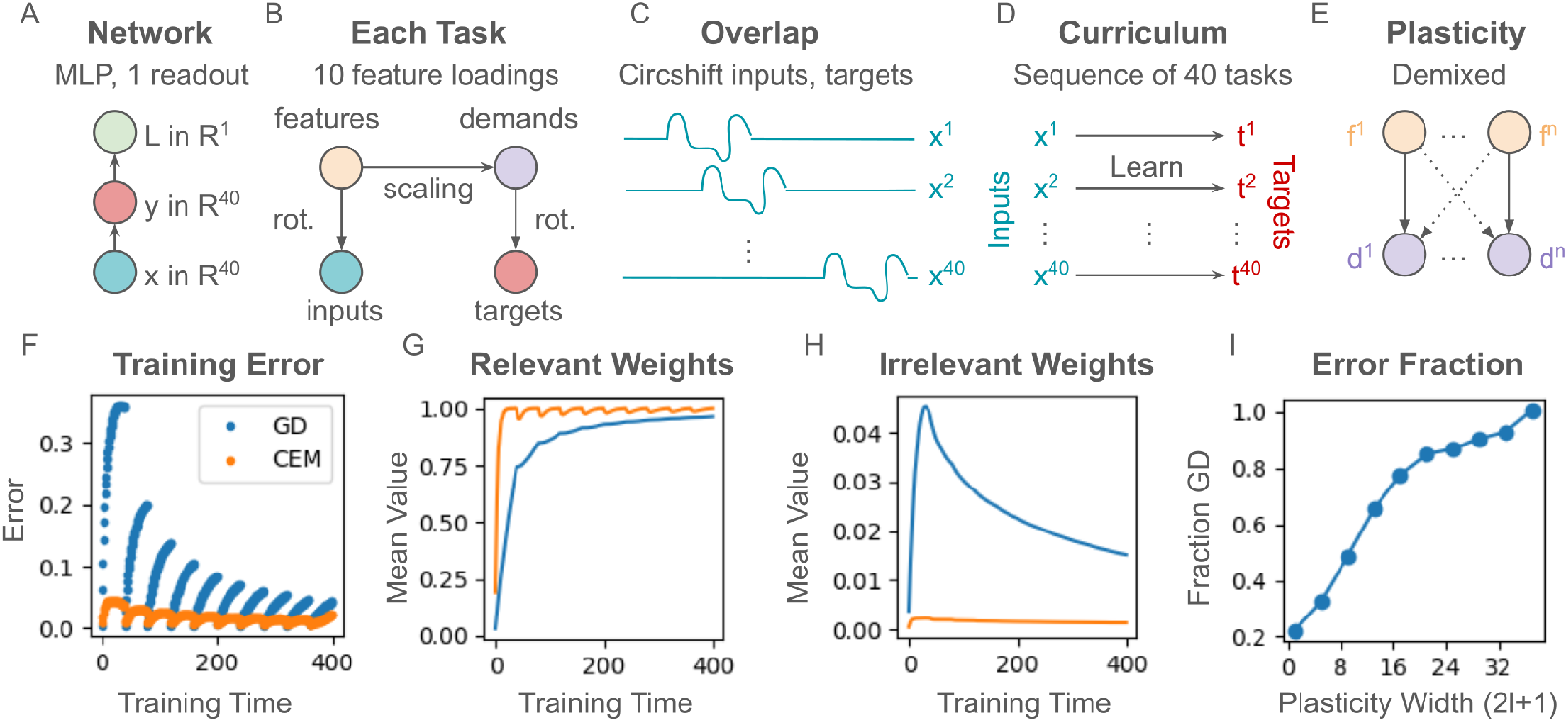
Compositional plasticity. (A) Networks were MLPs with single readouts. (B) Tasks for a curriculum were produced by taking 10 feature loadings, converting them via scaling to 10 task demands (latent outputs), and then generating observed inputs and targets as sums of features and demands (respectively). (C) Once input and target vectors were computed, we produced a curriculum of overlapping (interfering) tasks by circularly shifting them. (D) During training, networks learned to produce each target given each input. (E) Plasticity in the network was restricted to a sum of de-mixed subspaces (the representation in the network of each latent task demand could only learn about the representation of each feature). Unlike our other simulations, we did not perform unsupervised plasticity on gradient elements to first learn this de-mixing, as this learning problem is itself complex, and our main concern is the use of the PR-vs-RF eligibility decomposition itself. (F) Training error over 10 passes through the data, averaged over all tasks and over 50 simulation repetitions. The CEM converges to sub-criterion error with far fewer passes through the data than the network learning via GD. (G) Average weights between feature representations and their demand representations, with 1 being optimal. (H) Average weights between demands and features which are irrelevant for them, with 0 being optimal. Note that there are many more such spurious relationships than true ones. (I) Impact of linking number *l* (and hence dimensionality reduction) on cumulative training error over the 10-repetition window in F. Linking number 1 (corresponding to plasticity width 1 in the figure) represents completely accurate prior associations, whereas linking number greater than or equal to 20 (plasticity width 41) represents all-to-all plasticity between representations (gradient descent). The y-axis is cumulative error of the CEM as a fraction of GD.

**Figure 8.**
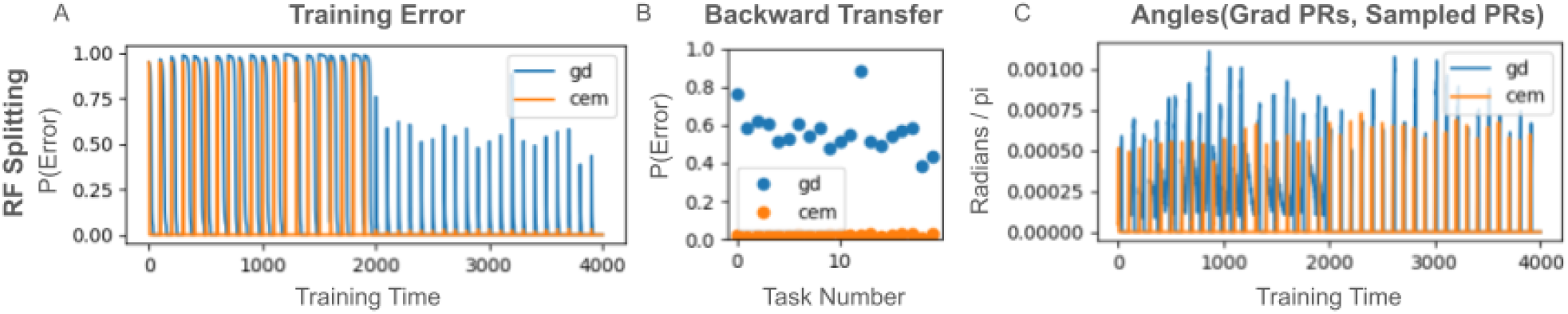
RF splitting in nonlinear neural networks that learn according to policy-gradient-like updates. (A) Training error for a simulation mirroring the RF splitting simulation (figure 3 from the main text). (B) The policy-gradient learner shows significant interference, whereas the coordinated eligibility model avoids this. (C). Sampled population response changes mirror those derived from analytic gradients in both models.

**Figure 9.**
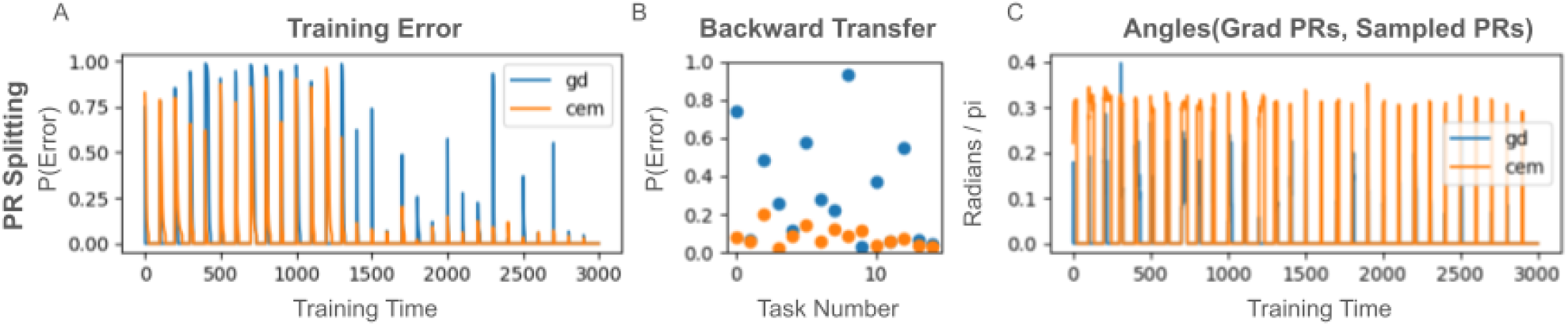
PR splitting in nonlinear neural networks that learn according to policy-gradient-like updates. (A) Training error for a simulation mirroring the PR splitting simulation (figure 4 from the main text). (B) The gradient-based method shows significant interference, whereas the coordinated eligibility model reduces this interference. Here, the CET does not totally abolish interference, because updates are only locally optimal, rather than globally optimal, owing to the curvature of the loss landscape induced by network nonlinearities. (C) Angles between CEM PR changes and gradient ones. CEM PR changes are consistently at a fairly high angle to analytic gradients, indicating that the locally optimal updates are nearly orthogonal to the gradient updates.

Our basic insight is that (unlike the earlier simulations) population response plasticity need not be independent of receptive field plasticity. Indeed, there is little reason to think these two aspects of response change are, in general, independent. Instead, as we consider here, subspaces of candidate population response changes (i.e., span({*u*_*i*_}), for some set *u*_*i*_) may be associated with subspaces of receptive field changes (span({ *v*_*j*_ }), for some set *v*_*j*_), leading to weight update matrices in span 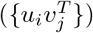, for *u*_*i*_ ∈ {*u*_*i*_} and *v*_*j*_∈{ *v*_*j*_}. When subspace associations such as these exist, the building blocks of representation changes are inherently yoked to those of receptive field changes. Learning multiple such updates simultaneously produces compositional learning, along with dimensionality reduction; instead of learning on the complete bipartite graph of possible population-response and receptive field updates, some smaller eligibility graph (with fewer edges) is thereby assumed.

As a biological example, striatal neurons multiplexing sensory information from different areas may have different population-level representations related to different sensory modalities, and it may be useful to explore holistic changes in these representations in a way that keeps learning somewhat separated by (for example, internal to) different modalities. This would imply yoking a representation change associated with one modality to a set of receptive field changes “listening” to that same modality. As noted above, we are not suggesting that other mechanisms cannot accomplish similar goals. Instead, we show that synaptic plasticity could be either pre-configured or tuned online to perform such functions in some circuits.

To explore this idea, we simulated simple, single-layer linear networks (figure 7A) solving compositional task sets. Tasks were constructed by generating random mathematical bases for network inputs and hidden layer responses, and associating elements of each basis with elements of the other. This defined how features (latent inputs, the input basis components), translated into latent task demands (the output basis components) in a 1-to-1 fashion (figure 7B). Since basis components for each feature were unit vectors, the inputs for each feature were generated as compositions of rotations and reflections (elements of the orthogonal group *O*(*n*)). The desired input-to-hidden transformation was thus an orthogonal matrix 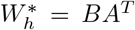, with input feature vectors encoded in *A* and task demands in *B* (figure 7B). Stimuli were generated as compositions of the basic features, with 10 random loadings per task, and target outputs were generated as compositions of graded responses to these input features (using the same loadings) (figure 7B). The strength of an input feature therefore determined the extent to which its associated output was required. Because there are 40-choose-10 features per task (which precludes using averages over draws to avoid introducing non-uniform sampling effects in our results), we selected stimuli to include all 40 features with equal frequency (figure 7C). Specifically, we did so by circularly shifting feature loadings to define a curriculum. These tasks were then learned sequentially (figure 7D), with plasticity in our coordinated eligibility model restricted to feature-demand subspaces which included the correct associations (figure 7E). We ran a series of such simulations, varying a “linking number” parameter, *l*, which determined the degree of plasticity restriction. This parameter varied between all-to-all (gradient) connectivity and 1-to-1 (completely accurate prior associations), and allowed us to examine the impact of restriction on learning.

Our results are illustrated in figure 7F-I, showing performance of the coordinated eligibility model relative to GD, averaged over 60 random task and network initializations. The coordinated eligibility model converges to its asymptotic error in fewer passes over the data relative to gradient descent, with less interference (figure 7F). This can be observed in the weights of the network, as relevant weights (defined as those connecting input features with their associated output demands) rapidly increase and plateau for the coordinated eligibility model, whereas gradient descent increases them much more slowly (figure 7G). Similarly, irrelevant weights (or spurious associations) grow much more rapidly for gradient descent, then decay, whereas the coordinated eligibility model suffers less from this problem (figure 7H) (the scales differ because there are many more irrelevant weights). Linking number impacted average cumulative error as expected, with even limited dimensionality reduction providing significant decreases in cumulative training error (figure 7I). Thus, feature matching via coordinated receptive field and response eligibility can drastically improve learning in compositional tasks relative to gradient descent.

## Discussion

Our results contribute to a tradition of work that shows how models of biological plasticity may have functions beyond those imparted by gradients. In this work, we showed that restricting plasticity to structured receptive field and population-response changes provides a powerful way to manage interference, and generalization. In managing generalization, we also illustrated links with the topic of state abstraction. Furthermore, by linking these constraints, networks can also learn compositionally. This compositional learning has the potential benefit of significantly reducing learning-problem dimensionality, among others. In support of these ideas, we introduced a plasticity equation encapsulating these different models. We called this umbrella description the coordinated eligibility theory, and showed how unsupervised learning on gradient components could produce the functions discussed above.

The plausibility of our model rests on the observation that modulatory neural inputs controlling plasticity are extremely numerous and diverse (reviewed in Scott & Frank 2022). In the cortex, for example, parvalbumin expressing interneurons such as basket and chandelier cells likely impact population response plasticity, via their impacts on local population-level firing rate variability. Similarly, somatostatin interneurons exert a degree of control over dendritic calcium concentrations, and (presumably) thereby receptive field plasticity (Higley 2014; Naka *et al*. 2019; Kecskés *et al*. 2020). Numerous other control mechanisms are also plausible, and as one would predict from our theory, experimental calcium manipulations in dendrites have already been shown to causally impact learning interference between tasks (Yang *et al*. 2014; Cichon & Gan 2015; Sehgal, Filho, *et al*. 2021).

The extent to which any given locus of plasticity displays either fixed or metaplastic gradient projections, along with what these projections are, would be expected to vary, and is a subject of further investigation suggested by our theory. Metaplasticity in action selection is most well established in the dopamine system, for example, (Yagishita *et al*. 2014; Collins & Frank 2014; Jaskir & Frank 2021; Scott & Frank 2022), where our theory suggests considering potential distinctions between D1 and D2 (dopamine receptor) pathways, which are associated with different aspects of cost and benefit calculations. In particular, these pathways specialize in decision-making under different circumstances (Collins & Frank 2014; Jaskir & Frank 2021), and they may further profit from considering how their PRs and RFs can be differentially sculpted (for example, to prevent unlearning in D1 MSNs while promoting new learning in D2 MSNs). As these examples suggest, our coordinated eligibility theory should useful in examining diverse circuits.

Our coordinated eligibility theory is also closely related to a number of other plasticity models, in both neuroscience and machine learning. As we discuss in the supplementary material, versions of the Bienenstock-Cooper-Munroe (Bienenstock *et al*. 1982), two-threshold calcium theories (Evans & Blackwell 2015), and neuromodulated Hebbian (or “three-factor”) rules (Frémaux & Gerstner 2016) are special cases of equation (4). As a result, our findings describe these models. Similarly, other continual learning schemes can be understood in terms of our results. For example, elastic weight consolidation (Kirkpatrick *et al*. 2017) can be considered an axis-aligned, less-structured version of our model, in the sense that weight values (and hence projections of weights onto axes in network weight-spaces) are fixed in ad-hoc non-decomposed ways. As further examples, memorization methods like gradient episodic memory (Lopez-Paz & Ranzato 2017) and replay-based methods (Wang *et al*. 2022) both essentially propose ways to remain within earlier tasks’ solution manifolds, which our networks do explicitly on the basis of gradient factors. By contrast with our work, however, these methods lack the the structural and functional insight provided by the decompositions and geometry we have discussed, and neither do they address generalization.

Nonetheless, our results here are limited in a variety of ways. Most notably, they can and should be extended to understand coordinated eligibility involving multiple non-linear network layers, and more complex combinations of learning subspaces. Here, we have focused on linear systems, because understanding phenomena in them is virtually always the foundation for understanding more complex cases. As such, our results here are applicable both infinitesimally within non-linear systems, as usual, and within linear network activity regimes induced by ReLU and GELU non-linearities, for example. They are therefore also applicable, in principle, to non-linear systems in general, but further work will be required to understand the details of such applications. In a similar manner, the subspace plasticity restrictions we discussed here can be considered as primitives in more complex settings. For example, it is likely that biological networks have a variety of more and less plastic subspaces within connected networks, such that plasticity is not a simple, binary function of connectivity. Our networks are the only ones we know of which describe this situation, and understanding biological plasticity will likely benefit from both such expanded understandings of coordinated eligibility models.

## Methods

### Network model

All simulations were carried out using PyTorch, running custom network layers. Code for these simulations can be found at www.github.com/DanielNScott/beyond-grads. Network layers implementing coordinated eligibility used forward and backward hooks to compute surprise as a function of inputs during network forward passes or as a function of supervision mismatch during backward passes. To do so, networks maintained histories of inputs and supervision, generally for one time-step of training, and compared new inputs or supervisory signals with those seen just prior. To compute gradient manipulations, layers also maintained state in the form of projection matrices. For pattern-separation simulations, these had previously seen dimensions removed, whereas for pattern-completion simulations, dimensions were merged by removing the independent (to-be-merged) dimensions and replacing them with one (joint) dimension. In our compositional plasticity simulation (simulation 5), the independent projections were not learned, being directly implemented instead.

### Surprise computations

During blocked learning, the appearance of a new task can be detected as a change-point in inputs or feedbacks, and likewise, these can be grouped over time according to contiguity. Empirical evidence suggests the locus coeruleus is heavily involved in signaling such (unvalenced) surprise (reviewed in Sara & Bouret 2012), whereas midbrain dopamine projections signal both valenced (reward-prediction error) and unvalenced surprise, associated with reinforcement learning and stimulus-stimulus association learning, respectively (Diederen & Fletcher 2021; Schultz *et al*. 1997). The neuromodulators these areas release, norepinephrine, and dopamine, have major impacts on plasticity (reviewed in Scott & Frank 2022). Our networks compute such surprise signals, and use them to perform pattern-separation (in the high-surprise case) or integration-based coarse-coding (in the low-surprise case) on plasticity components *u* and *v*, by changing the transformations *P* and *Q* over time.

Specifically, each layer computes a running average *µ* of its inputs over time. When new inputs arrive, surprise *s* is computed as the logistic function applied to cosine-angle dissimilarity between new and old inputs. That is:

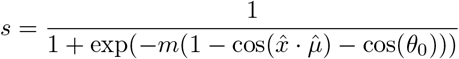

This surprise function is shown in figure 3E. The parameters *θ*_0_ and *m* define the shape (cutoff location and cutoff sharpness) of the sigmoid. The running mean computation is weighted by surprise, such that surprising events more fully replace the running average, and the transform *T*_*u*_ has the most-recent running mean removed according to surprise as well. That is, 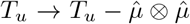 for a normalized mean 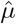. An analogous computation is performed on the *v* vectors, as we describe in the relevant section below. As we show below, these operations allow our networks to flexibly manage interference and generalization across tasks.

### Simulations 1 and 2

For each repetition of simulation 1, we initialized our network’s input eligibility transform as the identity matrix, *T*_*u*_ = *I*. After training on input *x*_*t*_ during the *t*-th task, *T*_*u*_ was updated as 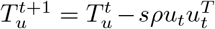, where *u*_*t*_ represents the vector 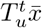 scaled to have unit norm, *ρ* is a mixture parameter, *s* is surprise (which was very close to 1 at task transitions, and 0 otherwise), and 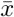 was the average input over the immediately preceding period of low-surprise (i.e., 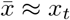, the input during the t-th task).

Plasticity in simulation 2 was treated analogously to plasticity in simulation 1, with the transform *T*_*v*_ being updated according to surprising output update supervision signals *g*_*t*_, rather than *T*_*u*_ being updated according to surprising inputs.

### Simulation 5

For simulation 5, we generated orthogonal bases *A* and *B* for the inputs and targets by randomly sampling multivariate normal distributions and applying the Gram-Schmidt procedure to orthogonalize them. The columns of *A* served as vectors mapping latent features to observed inputs, and the columns of *B* as vectors mapping latent demands to observed targets. We generated compositional input stimuli by circularly shifting (“rolling”) a fixed vector with *C* non-zero random weights *w*_*i*_, and using each of these as weights applied to the latent features. The same procedure was used to generate target outputs. For example, if stimuli are denoted *s*_*i*_, then in the *n* = 3, *C* = 2 case, we would have a set of inputs:

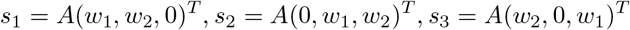

Juxtaposition here represents matrix multiplication, and parenthesis denote vectors (not indexing). Targets were generated using the same set of weights and the demand-target map *B* instead of the feature-input map *A*. Matching the inputs and targets shows that the learned optimal weight matrix will be 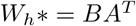, because this matrix first inverts the feature-to-input map (*A*) then applies to feature-to-demand map (*I*), and finally the demand-to-target map (*B*). This weight matrix was learned directly in the gradient descent simulations, and indirectly, in the latent feature-demand space, using the CEM. Since the basis transormations are arbitrary, both cases are equivalent to learning the identity map.

Linking numbers were free parameters that we used to encode prior information about the target weight transformation. In a simulation with linking number *l*, for each index *t*, plasticity was restricted to retain only terms between features [*t* − *l, t* + *l*] and their associated demands. For example, given a linking number of 1, plasticity would include terms 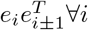 in the feature-demand space. A linking number of *l* = 0 therefore provided complete information about which features should be associated with which demands 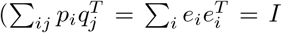, again, in the feature-demand space), such that only the strengths of these relationships required learning. A linking number such that 2*l* + 1 ≥ *n* was equivalent to gradient descent (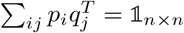, since *p*_*i*_ and *q*_*j*_ span all feature-demand basis pairs). These restrictions can be accomplished at by taking 2*l* + 1 projections of *g* and 2*l* + 1 projections of *x*, then summing all of their pairwise products, or they can be implemented parsimoniously using a mask *M*. The t-th row of this mask is then a vector of length *n* with 2*l* + 1 ones centered on the t-th component, with zeros elsewhere. We took the latter approach, transforming gradients into the feature-demand space, masking them, then transforming them back. Denoting the full gradient *G*, this meant that at each backward pass of the CEM network, we set *G*↦ *B*(*M* ⊙ *B*^*T*^ *GA*^*T*^)*A*.

## Acknowledgements

For helpful discussion, commentary, and feedback, we thank Matthew Nassar, Apoorva Bhandari, Rex Liu, Christopher I. Moore, Ian A. More, David Badre, Scott Susi, Cris Buc Calderon, and the Frank lab. Daniel Scott was supported by NIMH training grant T32MH115895 (PI’s:Frank, Badre, Moore). The project was supported by NIMH R01 MH084840-08A1. Computing hardware was supported by NIH Office of the Director grant S10OD025181.

## Author contributions

D.N.S. and M.J.F. developed the research topic. D.N.S. developed the mathematical analyses, functional and biological interpretations, wrote code, performed simulations, and drafted the manuscript. M.J.F. provided extensive feedback at all project stages and on all topics. D.N.S. and M.J.F redrafted and edited the manuscript, and prepared it for submission. D.N.S. and M.J.F revised it upon receiving feedback.

## Supplementary material

### Gradients, sampling, PRs, and RFs

In this section, we discuss some of the relationships between gradients, sample-based gradient estimates, and coordinated eligibility theories. To do so, we first show how to arrive at the PR-RF decomposition of the gradient, then we take the expectation (over noisy task repetitions) of a neuromodulated Hebbian rule to get a similar result.

Using Einstein notation, basis vectors *e*_*n*_, dual basis vectors *e*^*m*^, *h* as the element-wise derivative of *ϕ*, and *D*_*h*_ as matrix with *h* on the diagonal, the derivative of reward with respect to weights *W* is:

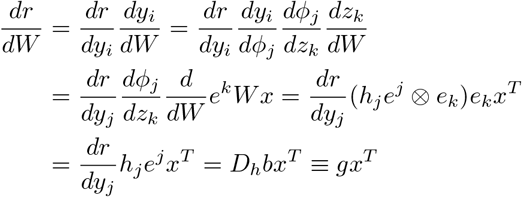

On the last line, we have defined *b* as (*dr/dy*)^*T*^ and *g*, the gradient output filter or gradient population response change, as *D*_*h*_*b*. This has the same form as our coordinated eligibility models.

The same form can be arrived at as the expected value of a set of reward-modulated Hebbian updates (Williams 1992). When these Hebbian updates are, on average, gradient-following, they are called REINFORCE algorithms (Williams 1992). Conceptually, it is important to note that REINFORCE algorithms are *defined* as gradient following methods, however, whereas neuromodulated plasticity is frequently defined mathematically, without respect to function (Frémaux & Gerstner 2016). As such, the two don’t always coincide. Moreover, even when a neuromodulated Hebbian rule is a REINFORCE algorithm for *some* network, that does not make it the REINFORCE algorithm for the *specific* network it is being applied to. With these points in mind, we can examine the following neuromodulated Hebbian rule, using a local linear expansion of *r* around 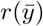, and denoting firing rate noise by *λ*_*y*_:

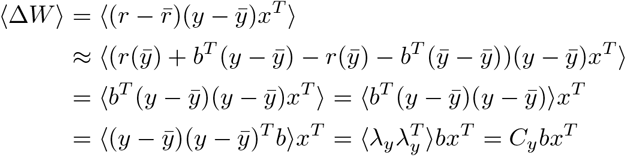

These equations indicate that when the output firing rate covariance *C*_*y*_ is proportional to the matrix of firing rate slopes *D*_*h*_, the neuromodulated Hebbian update is equal to the gradient in expectation. This specific conclusion is the same as that of Williams’ (1992) REINFORCE paper. More interesting, however, is the general reliance on *C*_*y*_, which was not calculated in that work. This term is often assumed to be isotropic (see, e.g., Fiete & Seung 2006), but in real neural networks, one would not expect it to be Kohn *et al*. 2016. In particular, when *D*_*h*_≈ *I*, which is (notably) always true for neurons in a GELU or ReLU network with firing rates reasonably far from zero, we have that *b* ≈ *g*, and hence *C*_*y*_ warps the gradient according to the main dimensions of variability in the network noise. In the case where *C*_*y*_ is not full rank, meaning that some dimensions have no noise, it projects those dimensions out of *g*. (Cases where the eigenvalue scales are very different interpolate between the projecting and non-projecting cases.) Since non-isotropic noise is more biologically plausible than isotropic noise, this provides another motivation for constructing the filter transforms *u*(*g*) and *v*(*x*) as we have in the main manuscript.

Finally, note that sampling can also be implemented upstream of a neuron’s activation function. Consider the update equation analogous to REINFORCE here, known as the node perturbation update (Fiete & Seung 2006):

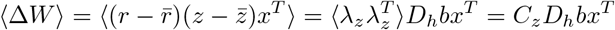

As with the previous equations, shrinking (i.e., regularizing) the covariance matrix here towards a low-rank matrix will project the gradient output filter.

### Coordinated eligibility results for reinforcement learning

The coordinated eligibility theory can be applied to supervised, unsupervised, or reinforcement-based plasticity. We applied it to supervised plasticity in the main text, without loss of generality, because this was the most straightforward demonstration to emphasize the geometric relationships between plasticity factors and their impacts on interference and generalization. Notably however, these results also apply to reinforcement-learning and unsupervised scenarios, which are connected to supervised learning through sampled and implicit gradients, respectively. Coordinated eligibility theories, by our definition, posit factor-structured transforms of these gradients, and do not require that such gradients are explicitly computed or represented.

In reinforcement learning, there are unique additional challenges due to the exploration exploitation problem, which impacts sample complexity. Network-based reinforcement learning requires searching over spaces of candidate actions, parameterized by network weights, in order to find good ones and propagate loss information. This makes the relevant search problems high-dimensional, and as a result, methods such as REINFORCE must accumulate many directional derivatives of gradients (i.e., sample many potential changes in their actions) in order to effectively follow loss gradients. These methods typically add noise to each neuron’s activity, and adjust synaptic weights via neuromodulated Hebbian plasticity. A well known limitation of such methods is that they require an inordinate number of samples to converge, relying on noisy perturbations to happen upon useful trajectories. Nevertheless, because they approximate gradient descent, these networks also inherit the usual deficiencies considered in the main text for the case of supervised learning, including catastrophic interference and lack of compositional generalization. Our Coordinated Eligibility Model can mitigate some of these deficiencies while remaining biologically plausible.

By restricting the number of dimensions involved in network plasticity, methods such as ours improve the sample complexity of RL methods, at the potential cost of moving in weight directions that have some significant non-zero angle relative to the un-restricted network gradients. To be clear, this cost is also the price paid for the benefit of avoiding interference, or promoting generalization, however. Previous work has shown how this sample complexity reduction can nonetheless improve learning outcomes (Nassar *et al*. 2021), but did not assess the impacts of different factor geometries, as we have here.

### Reinforcement learning simulations

In this section, we briefly describe simulations paralleling those in the main text, applying the CET to contextual bandit tasks. Contextual bandit tasks are simple RL scenarios in which states are visited randomly, and each state requires learning a single (stochastically optimal) response through trial and error. This is typically framed as visiting different casinos, wherein one has a set of slot machines to choose between playing; the context is the casino, the action is the slot machine play, the outcomes are in general stochastic, and the reinforcement learning problem is choosing the best machine in each casino based on reward receipt (which agents must essentially average over time). This setup is the reinforcement learning analogue of supervised input-output learning, such as labeling images. Contexts are analogous to images and actions to labels. The supervised case and the reinforcement one differ only in that actions must be tried and compared in order to determine the correct response for each input, with the additional caveat that feedback may be stochastic.

Any sensible RL model can solve contextual bandit tasks, but in networks, changing contexts generally induces experience-dependent forgetting, if learning is done online and in a blocked fashion. Furthermore, performing gradient descent on policies does nothing to manage generalization of learning from one context to another. These issues are inherited directly from relationships between gradients, as discussed in the main text. Issues related to exploration/sampling exacerbate them, but are fundamentally distinct. As such, coordinated eligibility models can control both interference and generalization in RL models, just as in the main text, which we demonstrate here.

Our reinforcement learning simulations used simple nonlinear three-layer neural networks, with 100 input neurons and 200 hidden neurons. RF-separation simulations used a single head with 10 output neurons, and PR-separation simulations used 20-unit heads for each task. Network weights were initialized according to log-normal distributions, while network outputs were softmaxed to get action probabilities, then subject to an expected-value loss. Neurons used softplus differentiable, positive-firing-rates. The networks were therefore non-linear, owing to both the softmax criteria and the softplus activations. During training, they learned sequences of 20 contextual bandit tasks, in a blocked fashion, with each block (single task training episode) containing 100 training trials. Backward transfer (destructive interference) was assessed by examining performance during a second epoch of training (a second training pass through the entire task set). To separate the CET’s impacts on sample complexity and on factor geometry, we defined “sub-trials” as loops of gradient accumulation steps. Each trial was composed of 500 sub-trials, and during each sub-trial, networks output an action probability distribution, and were presented with a reward-prediction error.

Our RF-splitting simulations (figure 1S) made use of 20 input-output pairs. Each input was a 40-hot binary vector, composed of a random portion and a deterministic one. The deterministic component was used to guarantee every vector had at least one input that was unique, whereas the random component provided overlap. Specifically, the unique portion of each input was a 1-hot vector of length 20, which was prepended to a randomly permuted 40-hot vector of length 80. Each input had a unique associated target action (1-hot of length 20 serving as a softmax output target). Networks were trained to produce these actions as described above, using either a policy-gradient method (REINFORCE) or a projected policy gradient (our CET). Projections were defined offline to isolate the unique elements of different inputs and project policy gradient input filters onto these components. (Note that this can also be done online, as in the main text). Training curves over the whole simulation (figure 1S, A) illustrate that both methods completely learn each task. The policy gradient method produces significant forgetting (figure 1S, A and B), however, whereas the coordinated eligibilty model does not. Finally, the accuracy of the sampled population response filters can be seen to be equivalent across the two simulations (as it should be, figure 1S, panel C).

Our PR-splitting simulations (figure 2S) also consisted of 20 input-output pairs. Like the tasks in the text, these all used the same constant input, and each required a single target action. Each task used a different readout head, which was a random matrix with i.i.d. log-normal entries, randomly scaled by row (fixed across epochs) to have mean 1-norms of 1, with a standard deviation of 0.2. Hidden weights were distributed according to the same log-normal distribution parameters. PR projections were defined with each task’s firing-rate sample space in the joint null-space of all other task’s readout weights. These projections were pulled back to the population response space on z-values through the instantaneous firing rate nonlinearities, for sampling. As above, both policy gradient and coordinated eligibility networks learned the all tasks completely (figure 2S, panel A), while the policy gradient updates produced significant backward transfer (figure 2S, A and B). Unlike the RF simulation, population response changes had consistent angles of roughly 0.4*π* relative to true gradients (figure 2S, C), indicating that the optimal updates (for the whole curriculum) were very nearly orthogonal to the gradient updates.

Analogous effects to those seen in figures 5,6, and 7, relating to generalization and composition of plasticity components can also be shown in reinforcement learning simulations.

